# CDH-3/Cadherin, YAP-1/YAP and EGL-44/TEAD promote SYX-2/Syntaxin and EFF-1 fusogen-mediated phagosome closure

**DOI:** 10.1101/2025.04.02.646655

**Authors:** Alec Whited, Aladin Elkhalil, Ginger Clark, Piya Ghose

## Abstract

Physical interactions between cells, such as cell-cell junctions, can profoundly impact cell fate. A vital cell fate for normal development and homeostasis is programmed cell death. Cells fated to die must be efficiently cleared away via phagocytosis, and defects are associated with a variety of diseased states. Whether cell-cell physical associations affect programmed cell elimination has not been well-explored. Here we describe, *in vivo,* a cell-cell adhesion-driven signaling pathway that ensures compartment-specific cell clearance during development. We previously described the specialized cell death program “Compartmentalized Cell Elimination” (CCE) in the *C. elegans* embryo. During CCE, the tail-spike cell (TSC), a polarized epithelial cell, undergoes a tripartite, ordered, and organized death sequence, allowing for the study of three distinct death modalities in a single cell setting. Prior to its demise, the TSC serves as a scaffold for the tail tip, formed by the hyp10 epithelial cell which develops along the TSC process. The hyp10 cell in turn also serves as the phagocyte for the dying TSC process. Here we present data suggesting that the physical association between the dying TSC and hyp10 phagocyte via CDH-3/cadherin mediates function of the mechanosensitive transcriptional coactivator YAP-1/YAP and its partner EGL-44/TEAD in the hyp10 phagocyte to promote localization of hyp10 SYX-2/Syntaxin around the dying TSC remnant. This pathway facilitates the phagocytic function of EFF-1/fusogen, which we have previously shown to be required for phagosome sealing during CCE. Our work sheds additional light on a poorly understood step of phagocytosis and implicates adhesive forces and signaling between cells as important in cell uptake.

## Introduction

Cells encounter physical force as a part of their native environment and this serves as an important interaction that influences cell fate, behavior, function, and maintenance (1) (2) (3). Cells can encounter physical forces from neighboring cells. Cell-cell adhesion machineries are key in mediating the physical communication between neighboring cells (4). Defects in cell-cell junctions, where these adhesion machineries function, are linked to several disease states, including cancer (4). Adherens junctions are one type of cell-cell junction that can remodel to control dynamic cellular behaviors (5).

Cadherins are Ca^2+^-dependent adhesion molecules that function in regulating epithelial cell morphogenesis (6). Cadherins have been linked to cell elimination in various contexts, including apoptosis (7) and entosis (8). Interestingly, E-cadherin-mediated cell-cell adhesion is involved in the coordination of neighbor cell elongation during apoptotic extrusion (9). Cadherins are also molecularly linked to phagocytosis, as in the case of Cadherin-11 (CDH11) of macrophages (10).

YAP (YES-associated protein) is a transcription regulatory factor that has several roles in general development, such as cell proliferation (11). Defects in YAP or its regulation have been implicated in cancer (12). YAP has also been identified as a conserved mechanotransducer that can translate various mechanical cues to defined transcriptional programs (13). YAP has been shown to act in tumor cells to allow them to avoid phagocytosis by macrophages (14). In response to cell fusion, YAP has been shown to exit the nucleus (15). The *C. elegans* homolog of YAP, YAP-1 has been shown to have several roles including in pharynx size scaling (16), thermotolerance and aging (17). and neuronal polarization (18).

Cells that undergo programmed death must also be efficiently cleared (19). Programmed cell death is most extensively characterized in the case of apoptosis, and phagocytosis of apoptotic cells is well-studied (20). Phagocytosis of non-apoptotic cells has also been studied (21). Phagocytosis is a process that allows for the clearance of numerous types of cargo such as dead cells, cell debris, and microbes by cellular ingestion and is vital to tissue homeostasis, immunity, and development. Studies have revealed the transcriptional signatures of different types of phagocytes, such as macrophages (22) and microglia during phagocytosis (23). This process involves dramatic yet controlled reorganization of cell morphology around the cargo that is limited by physical constraints involving relevant aspects of the membranes, receptors, and cytoskeleton (24). Phagocytosis entails a series of defined steps, beginning with corpse recognition, followed by extension of the phagocyte pseudopods to begin enveloping the corpse. The protrusions of this advancing phagocytic cup reach a point of contact and merge. The nascent phagosome separates from the plasma membrane via membrane fission. Sealing of these pseudopods around the corpse to form the phagosome vesicle is then followed by a series of phagosome maturation events (25, 26), culminating in fusion of lysosomes to the phagosome that contribute hydrolases to complete the corpse degradation process. Most of these steps are well-described. For example, apoptotic corpse recognition is well-known to be motivated by the presentation of phosphatidyl serine (PS) on the corpse membrane as an “eat-me” signal (27). Phagosome closure is the final critical step of cargo internalization but arguably most poorly understood (28). It has been postulated that one reason why phagosome sealing is a challenge to study is the difficulty in actually distinguishing fully enveloping but unclosed phagosomes (24). For phagocytosis of polarized cells, such as neurons, an important consideration is the compartmentalized nature of such cells and that different regions of these cells may be killed and cleared differently. Stereotyped pruning is a fundamental feature of nervous system development (29). Morphologically, pruning can be divided into two types. In the first type, axons become fragmented as pruning proceeds, in a manner similar to axon degeneration. An example is Drosophila mushroom body (MB) axon remodeling (30) and mammalian visual cortical spinal tract projection removal (31) . The other form of pruning is retraction-like. Here axons draw back and do not fragment, for example, in rodent hippocampal infrapyramidal tract (IPT) during postnatal development (32). Notably, the phagocytosis of axonal fragments by glial cells is well known (29). In *C. elegans,* glia play a role in sensory neuron sculpting via phagocytosis (33). The relationship between retraction and phagocytosis is not well-studied.

Plasma membrane fusion is an important developmental event (34). There is evidence of mechanical force promoting cell fusion (35) and mechanosensory responses occurring at fusogenic synapses (36). EFF-1 is a transmembrane protein shown to act in homotypic cell-cell fusion during *C. elegans* development as well as when expressed in insect cells (37). Its ability to act as a fusogen is a function of its dynamic localization at the fusion-fated borders of the fusing cell (38). EFF-1 is important for *C. elegans* seam cell stem-like fate (39, 40), as well as membrane repair (41). EFF-1 cell surface exposure has been shown to be regulated by SYX-2 (41), RAB-5, and DYN-1. SYX-2/Syntaxin2 has been shown to interact with the EFF-1 C terminus to promote EFF-1 recruitment to membrane fusion sites. SYX-2 is a type of soluble N-ethylmaleimide-sensitive factor attachment protein receptors (SNARE) protein. SNAREs are key molecules for eukaryotic membrane fusion (42). DYN-1/Dynamin and RAB-5 remove EFF-1 from the plasma membrane of epidermal cells by endocytosis which then accumulate at early endosomes. Cell-cell fusion occurs only at membranes where EFF-1 is transiently and dynamically localized (43). We have previously described the tripartite specialized cell death program Compartmentalized Cell Elimination (CCE) in *C. elegans* in both the tail-spike epithelial cell (TSC) and sex-specific sensory CEM neurons (44). CCE is defined by a highly ordered and stereotyped cell elimination sequence with three regression morphologies visible in three different compartments of the same specialized cell. The TSC is a polarized cell with a long tail-directed process (**Supplemental Figure S1a, intact stage**). The cell soma rounds, the proximal segment of the single process/dendrite undergoes beading and fragmentation (**Supplemental Figure S1b, beading stage, BA, S1c, beading not attached, BNA**), and the distal segment retracts into itself (**Supplemental Figure S1d, soma distal retracting stage, SDR**) leaving behind a regressed soma corpse and process remnant which are subsequently phagocytosed by different phagocytes stochastically (**Supplemental Figure S1e soma-distal degrading, SDD, early, Supplemental Figure S1f, soma distal degrading, SDD, late**). TSC process retraction is reminiscent of the forms of pruning. TSC death has been shown to be dependent on the main *C. elegans* caspase, CED-3, but not on its upstream regulator EGL-1/BH3-only (45). Studies on the TSC have identified multiple non-apoptotic players in programmed cell death (45–47). We have also previously shown that phagocytosis of the different compartments of the TSC involves distinct programs (44). The TSC soma requires the canonical engulfment pathway involving CED-5/DOCK180 for its engulfment, whereas the process does not. Clearance of the process on the other hand requires EFF-1 fusogen, which seals the phagosome formed around the distal process remnant. EFF-1 does not appear to play a role in soma phagocytosis and is in fact not expressed in the soma-neighboring phagocyte. Earlier studies on electron micrographs suggest that the TSC acts as a scaffold for tail formation (48).

Here we propose that, as CCE progresses, the resulting physical interaction between the TSC distal node and the surrounding hyp10 phagocyte via CDH-3/Cadherin activates the mechanosensitive YAP-1/YAP and EGL-44/TEAD transcription factors. This helps the SNARE protein SYX-2/Syntaxin of hyp10 localize around the diminishing TSC process to in turn promote the translocation of EFF-1/fusogen to hyp10 pseudopods to complete phagosome closure and corpse internalization. Our study highlights how the physical association between neighboring cells can guide cell elimination, and introduces a transcriptional control axis for phagosomal sealing, a poorly understood step of phagocytosis.

## Results

### CDH-3/cadherin promotes TSC corpse internalization

The TSC serves as a physical scaffold to the developing hyp10 tail-tip cell prior to CCE (44). We asked whether genes involved in the cell-cell physical association between these cells affect CCE and performed a candidate gene screen. We first considered cadherins. There are 12 *C. elegans* genes that encode cadherins (49). Of these, we noted from prior literature that mutants for the gene encoding the fat-like protocadherin, CDH-3, have hypodermal cell and tail-tip defects (50). Upon examination of a *cdh-3* transcriptional reporter (*cdh*-3 promoter-driven mKate2), we found expression in both the TSC and hyp10 (**Figure 1a**), as would be expected of a role of cadherins in cell-cell association. We proceeded to examine our previously employed TSC membrane reporter in two *cdh-3* mutant backgrounds (*pk87, pk77,*(*50*)) (**Figure 1b, c)**. Interestingly, we did indeed find CCE defects in these mutants, a range, predominantly distal process remnants. While the reported and observed L1 larval tail-tip morphology defect ((50) and **Figure 1b**) is seen 100% of the time, the CCE defect is not, suggesting these two phenotypes are separable.

**Figure 1.**
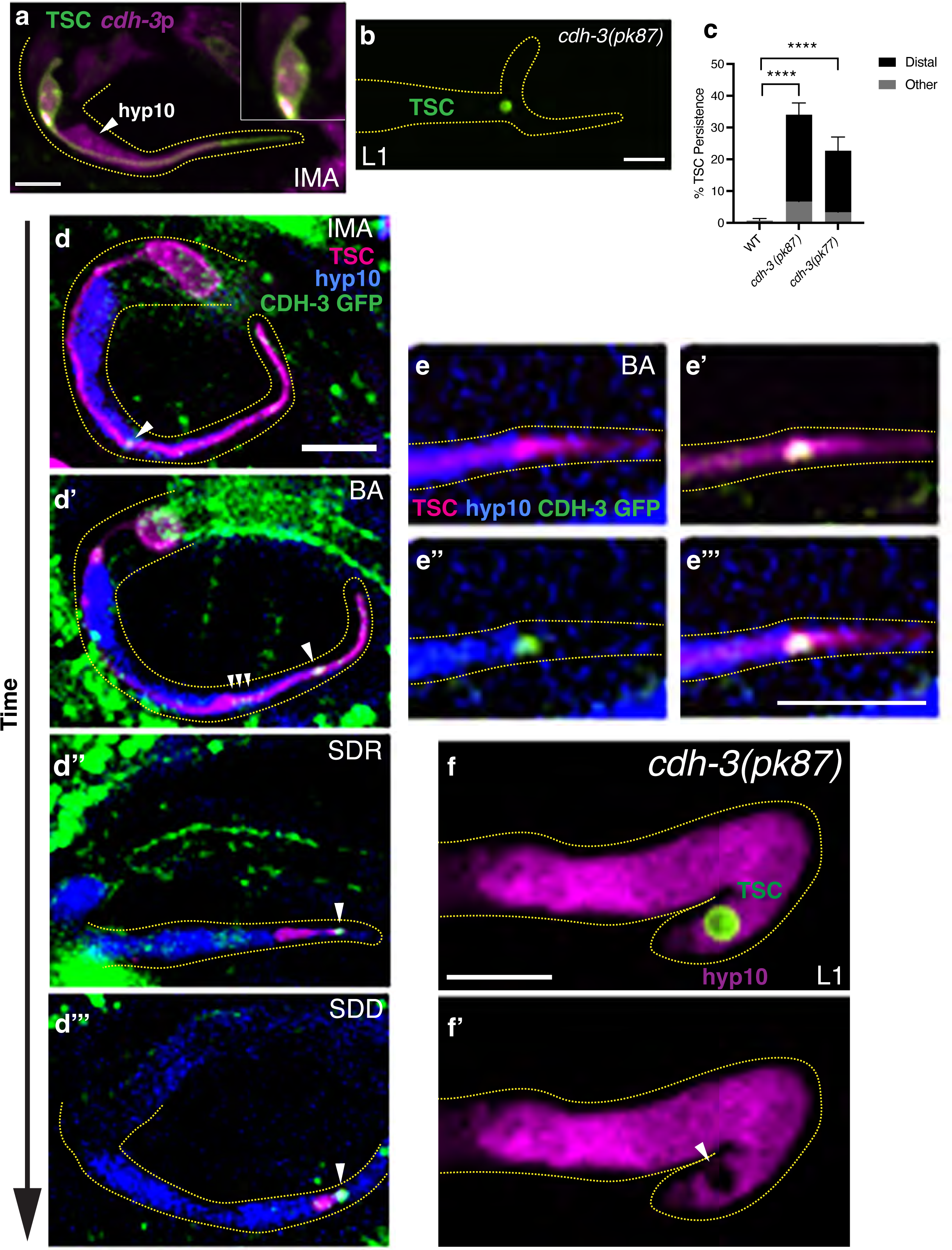
CDH-3/cadherin promotes TSC corpse internalization. **(a)** *cdh-3* expression in both TSC and hyp10. (b) CCE defect in L1 of *cdh-3(pk87)*. (c) Graph of *cdh-3* mutant allele phenotypes *cdh-3(pk87)* and *cdh-3(pk77)*. (d-d’’’) CDH-3::GFP protein localization across TSC development and CCE stages, showing adjacent presence in both TSC and hyp10 with white arrowhead showing CDH-3 localization. (e-e’’’) Tricolor labeling of CDH-3 (GFP), TSC (mKate2) and hyp10 (iBlueberry showing association of CDH-3 with both TSC and hyp10 Representative localizations 5/5 for each CCE stage. (f, f’) Test for *cdh-3(pk87)* mutant TSC remnant internalization with white arrowhead showing phagocyte opening. Scale bar: 5μm. For images of *cdh-3* expression in TSC and hyp10 n=10. For mutant scoring n>50. For *cdh-3(pk87)* internalization n>10. ns (not significant) *p* > 0.05, * *p* ≤ 0.05, ** *p* ≤ 0.01, *** *p* ≤ 0.001, **** *p* ≤ 0.0001.

We next sought to examine the localization of the CDH-3 protein across different CCE stages by employing a CRISPR/Cas9 generated strain in which GFP was introduced into the endogenous locus of the *cdh-3* gene at the C-terminus. We created this CRISPR/Cas9 strain in the previously employed background of TSC promoter-driven membrane mKate2 and hyp10 promoter-driven iBlueberry (44) (**FIG 1d-1d’’’)**. Examining this tricolor reporter, we found that CDH-3 distinctly localized in both the mature TSC (**Figure 1d**) and hyp10 at the TSC and hyp10 junction of the TSC distal process. This becomes more prominent at the CCE beading stage with some accumulation around the distal node formed in the TSC (**Figure 1d’**) as the distal part of the process retracts, with more prominent accumulation posteriorly (**Figure 1d, e**). Based on this defined spatiotemporal pattern and the mutant CCE defects, we proposed that CDH-3 and cell-cell adhesion are important for CCE. Unexpectedly, we also noticed that the hyp10 appears to begin to recognize the distal node at the beading stage forming what appears to be a phagocytic cup (**Supplemental Figure S2**). This leads us to speculate that the phagocytic process may begin at this stage, and that perhaps the change the adhesion and mechanical relationship between the TSC and hyp10 in the distal node region serves as a non-canonical “eat-me signal”. Indeed, we previously reported that the TSC process, unlike the soma, is not engulfed via the canonical apoptotic clearance pathway (44, 50). This potentially novel corpse (or future corpse) recognition mechanism will be an interesting future study.

We then asked whether the TSC remnants seen in *cdh-3* mutants at the L1 larval stage (which are not seen in wild-type at this stage) have been internalized by the hyp10 phagocyte. We used a cytosolic reporter for hyp10 (*skn-1*p::mKate2) and found that, while pseudopods appear to form, the TSC distal process remnants are not fully internalized (**Figure 1f, f’**). This suggests the TSC remnants of *cdh-3* mutants are still recognized by the hyp10, but that there is a failure in the actual internalization, suggesting a phagosome closure defect.

### CDH-3/Cadherin promotes EFF-1/Fusogen localization to nascent phagosome

We next asked whether the corpse internalization defect we see in *cdh-3* mutants is one of phagosome closure failure. We have previously reported that the cell-cell fusogen EFF-1 promotes phagosome sealing during CCE (44). In *eff-1* mutants as well, the distal process of the TSC persists and the hyp10 phagosome remains open resulting in a failure to internalize the TSC (44) (**Figure 2a, b, b’**). Here too, in support of our previously reported evidence, we showed that in *eff-1* mutants the tail morphogenesis defect and the CCE defect are separable (100% tail tip defects at L1) (44). We asked whether the CCE defects of *cdh-3* mutants and *eff-1* mutants are linked. We observed that the *cdh-3* and *eff-1* mutant CCE defects phenocopy and are not additive (**Figure 2c**). We then overexpressed *eff-1* in hyp10 expressing *eff-1* previously used to rescue the *eff-1* mutant phenotype (44) in *cdh-3(pk87)* mutants and found this to rescue the *cdh-3* defect as well (**Figure 2d**). These data suggest that EFF-1 functions downstream of CDH-3 to promote phagosome sealing during CCE. As we have previously shown, we report here using a tri-color reporter for the TSC (mKate2), hyp10 (iBlueberry) and EFF-1 (GFP) that EFF-1 localizes to the phagosome pseudopods (44) (**Figure 2e-e’’’**). To test the idea that CDH-3 may regulate this localization of EFF-1, we examined this tri-color line in the *cdh-3(pk87)* mutant background. Interestingly, we found that EFF-1 failed to localize/localize properly to the hyp10 phagocyte pseudopods in the *cdh-3(pk87)* mutant (**Figure 2f-g’’’**), though it is still seen on the membrane elsewhere. This suggests that CDH-3 promotes the translocation of EFF-1 to the phagocytic pseudopod. We therefore identify CDH-3 as a new regulator of phagosome sealing, a poorly understood step of cell clearance and a positive regulator of EFF-1 localization to the phagosome.

**Figure 2.**
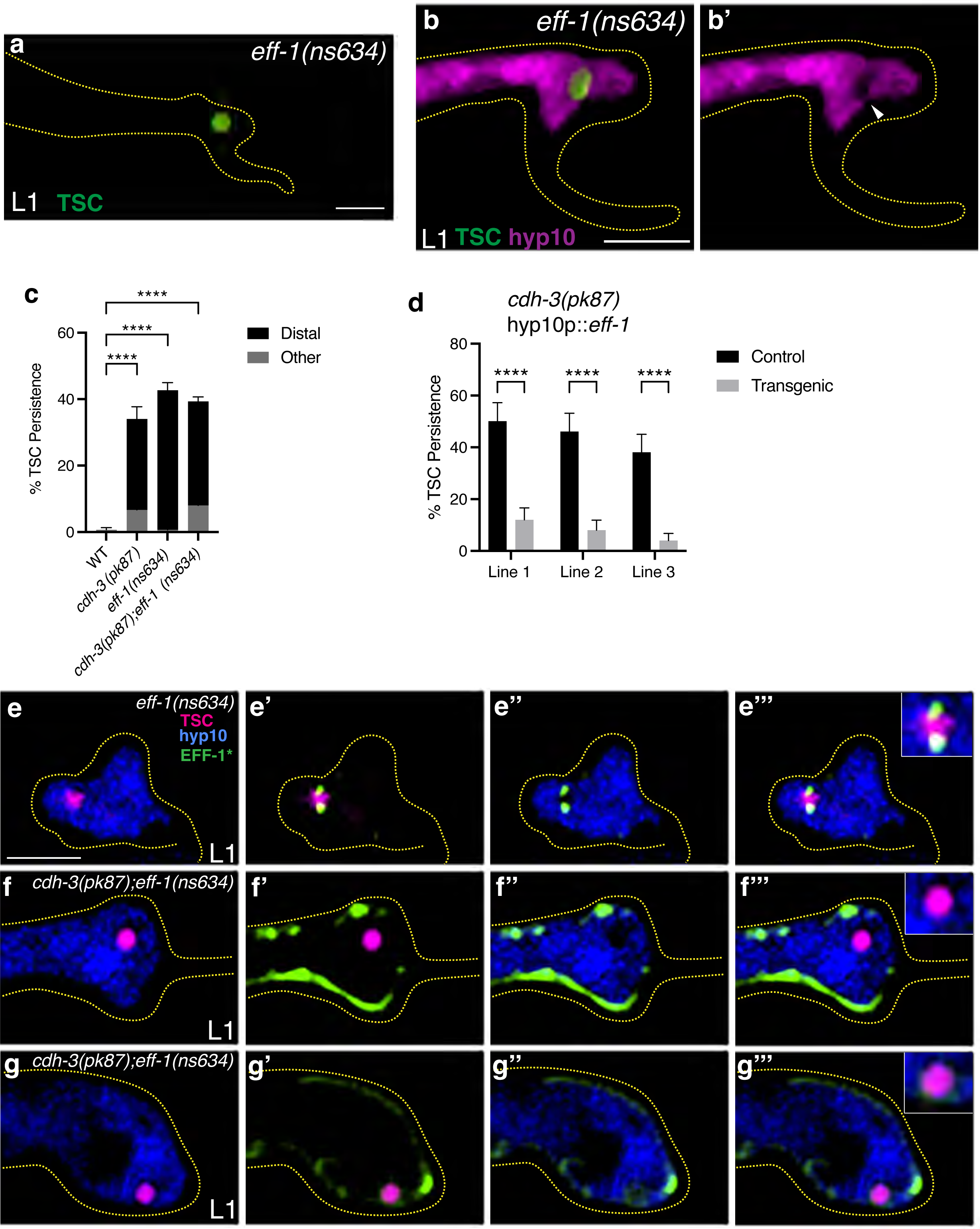
CDH-3/Cadherin promotes EFF-1/Fusogen localization to nascent phagosome. (a) CCE defect in L1 of *eff-1(ns634)*. (b, b’) Test for *eff-1(ns634)* mutant TSC remnant internalization with white arrowhead showing phagocyte opening. (c) Graph comparing *cdh-3(pk87)*, *eff-1(ns634)* single mutant and combination *cdh-3(pk87);eff-1(ns634)* defects. (d) Graph CCE defects for hyp10 overexpression of *eff-1(ns634)* in *cdh-3(pk87)* mutants. (e-g’’’) Tricolor labeling of EFF-1 (GFP), TSC (mKate2) and hyp10 (iBlueberry) at L1 larval stage in wild-type and *cdh-3(pk87).* Scale bar: 5μm. For over expression lines n>50. For mutant scoring n>50. For *eff-1(ns634)* internalization n>10. ns (not significant) *p* > 0.05, * *p* ≤ 0.05, ** *p* ≤ 0.01, *** *p* ≤ 0.001, **** *p* ≤ 0.0001.

### CDH-3/Cadherin promotes phagosome closure via hyp10 phagocyte YAP-1/YAP and EGL-44/TEAD

We next asked how mechanistically CDH-3/cadherin, by definition involved in the communication between cells, could promote successful TSC internalization via hyp10 EFF-1-mediated phagosome sealing. To address this, we looked into molecularly linking cell-cell adhesion and phagosome closure. We considered the fact that cell-cell adhesion entails a physical communication between cells and the fact the CDH-3 appears to localize, and presumably function, once the TSC distal node forms, and the dying cell and its phagocyte become more closely associated.

With this idea in mind, we thought of molecules that have been linked to responses to mechanical force. Previous work in cultured cells has shown that mechanical strain can induce E-cadherin-dependent activation of mechanosensitive transcriptional co-activator Yap during cell cycle entry (51), though to our knowledge a role for YAP in cell elimination has not been reported. We considered the *C. elegans* YAP homolog, YAP-1 (52). We examined mutants for *yap-1(ys38)* and found these to phenocopy both *cdh-3(pk87)* and *eff-1(ns634)* mutants (**Figure 3a, c-c’, e**). We also found that the *yap-1(ys38)* mutant defect could not be rescued via TSC promoter-driven *yap-1* cDNA (**Figure 3f**). However, we found that the *yap-1(ys38)* mutant defect rescued when expressing *yap-1* in hyp10 (**Figure 3f’**). This suggests that, as we have shown for *eff-1* mutants (44), YAP-1 functions non-autonomously in the hyp10 cell to promote CCE. We next tested whether the TSC remnant of *yap-1(ys38)* mutants has been internalized by hyp10, using the internalization reporter above. Here too we found, as for *cdh-3(pk87)* and *eff-1(ns634)* mutants, that hyp10 pseudopods were not fully closed around the TSC remnant (**Figure c, c’**), suggesting that, like CDH-3 and EFF-1, YAP-1 acts in hyp10 to promote phagosome sealing, perhaps molecularly linking the cadherin and the fusogen. To explore this idea further, we looked at *yap-1(ys38);cdh-3(pk87)* double mutants and found that the defects were not additive (**Figure 3e**).

**Figure 3.**
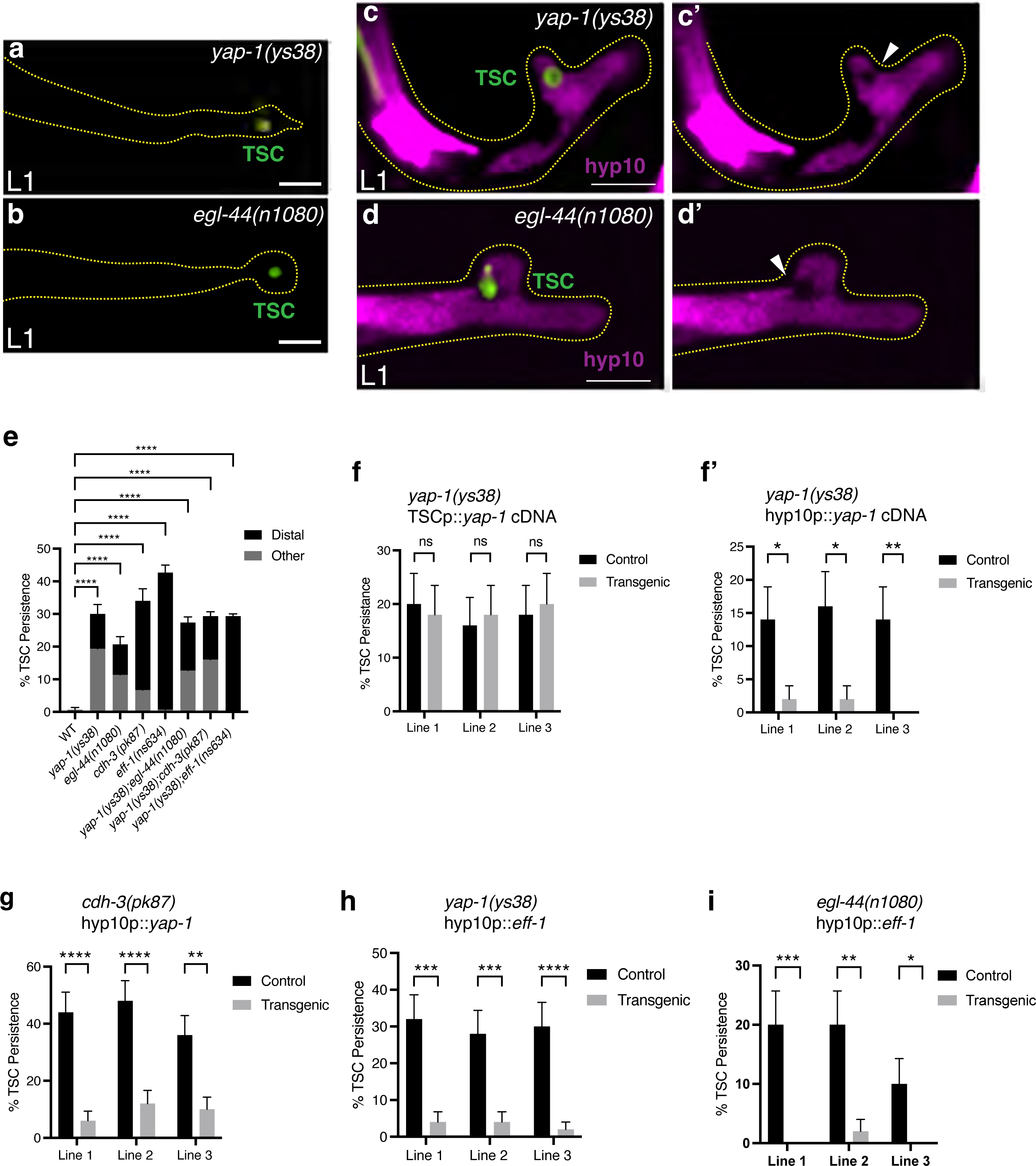
CDH-3/Cadherin promotes hyp10 phagocyte YAP-1/YAP and EGL-44/TEAD during phagosome closure. CCE defect in L1 of *yap-1(ys38)* **(a)** and *egl-44(n1080)* **(b). (e)** Graph quantifying *yap-1(ys38), egl-44(n1080) and cdh-3(pk87)* single mutant and combination *yap-1(ys38);egl-44(n1080)*, *yap-1(ys38);cdh-3(pk87)*, *yap-1(ys38);eff-1(ns634)* defects. Test for TSC remnant internalization in *yap-1(ys38)* **(c, c’**) and *egl-44(n1080)* **(d, d’**) with white arrowhead showing phagocyte opening. Graph of TSC **(f**) and hyp10 **(f’**) *yap-1* cell specific rescue. (g) Graph of *yap-1* overexpression in *cdh-3(pk87)* mutants. **(h)** Graph of hyp10 *eff-1* overexpression in *yap-1(ys38)* mutants. **(i)** Graph of hyp10 *eff-1* overexpression in *egl-44(n1080)* mutants. Scale bar: 5μm. For over expression lines n>50. For mutant scoring n>50. For *yap-1(ys38)* internalization n>10. For *egl-44(n1080)* internalization n=5. ns (not significant) *p* > 0.05, * *p* ≤ 0.05, ** *p* ≤ 0.01, *** *p* ≤ 0.001, **** *p* ≤ 0.0001.

Similarly, we found no additive effect of loss of *yap-1(ys38)*;*eff-1(ns634)* (**Figure 3e**). We further overexpressed the hyp10 specific *yap-1* rescue construct in *cdh-3(pk87)* mutants and found rescue of the *cdh-3* mutant defect (**Figure 3g**). We then overexpressed hyp10-specific *eff-1* in *yap-1(ys38)* mutants and found rescue of the *yap-1(ys38)* mutant defect (**Figure 3h**). This supports the idea that YAP-1 acts downstream of CDH-3 to promote EFF-1-dependent phagosome sealing during CCE.

We then tested for an involvement of EGL-44 (52), the worm homolog of TEADs (TEA DNA-binding domain proteins). TEADs are known to function as the main transcription factor partners of YAP (53). In this instance, we did find that *egl-44(n1080)* mutants had a similar CCE defect as *yap-1(ys38), cdh-3(pk87)* and *eff-1(ns634)* mutants (**Figure 3b, d, d’, e**). We also found that loss of both *egl-44(n1080)* and *yap-1(ys38)* did not have an additive CCE defect (**Figure 3e**), suggesting EGL-44/TEAD functions in the same pathway as YAP-1/YAP to promote CCE. We next tested whether the TSC remnant of *egl-44(n1080)* mutants has also been internalized by hyp10, using the internalization reporter above. Here too we found that hyp10 pseudopods were not fully closed around the TSC remnant (**Figure d, d’**), suggesting that EGL-44 also acts in hyp10 to promote phagosome sealing. We then overexpressed hyp10-specific *eff-1* in *egl-44(n1080)* mutants and found rescue of the *egl-44(n1080)* mutant defect (**Figure 3i**). This supports the idea that, like YAP-1, EGL-44 acts downstream of CDH-3 to promote phagosome sealing during CCE by facilitating EFF-1 function.

### CDH-3, YAP-1 and EGL-44 are important for defined localization of SYX-2/Syntaxin at TSC during CCE

We further wanted to probe how CDH-3, YAP-1 and EGL-44 direct EFF-1-mediated phagosome sealing and specifically looked for candidate transcriptional targets for EGL-44/TEAD that may also be involved in EFF-1 regulation. The SNARE protein SYX-2/Syntaxin 2 has been identified as a regulator of EFF-1 translocation to fusion membranes in the context of wound healing in *C. elegans* (41), Moreover, mammalian syntaxins are predicted to be transcriptional targets of TEAD1 (54, 55).

With these points in mind, we tested the hypothesis that the *syx-2* gene is the target of YAP-1/YAP EGL-44/TEAD as part of a CDH-3/cadherin-mediated signaling pathway in response to physical association changes between the TSC and hyp10. To this end we generated an endogenous reporter for SYX-2 using CRISPR/Cas9 by introducing GFP into the N-terminus of the endogenous locus of the *syx-2* gene in our strain in which the TSC membrane is labelled with mKate2 and hyp10 with iBlueberry. We then examined SYX-2 expression and localization across CCE stages relative to the TSC and hyp10. We first validated this newly generated SYX-2 reporter. Previous studies have shown that SYX-2 accumulates at wound sites (41). We similarly injured the hypodermis of adult animals using a microinjection needle and found SYX-2 of our CRISPR strain to prominently localize at injury sites (**Supplemental Figure S3**). This gave us confidence that we would be able to visualize SYX-2 during CCE.

We proceeded to examine the expression and dynamics of SYX-2 in the context of the TSC and hyp10 during CCE using this tricolor reporter **(Figure 4a-e’**). We found that SYX-2 signal can be first seen in hyp10, but not the TSC, from the CCE beading stage. The SYX-2 signal becomes more enriched at the center of the regressing distal process (**Figure 4c, c’ and d,d’**). Interestingly, SYX-2 is enriched in hyp10 at a region that appears to be wrapping around the center of bi-lobed distal process of the CCE SDD stage. This leads us to speculate that hyp10 SYX-2 may play a role in non-autonomous scission of the TSC distal process enroute to clearance. Alternatively, given its canonical role of membrane fusion, SYX-2 may be involved in hyp10 membrane autofusion as a part of the TSC distal process engulfment.

**Figure 4.**
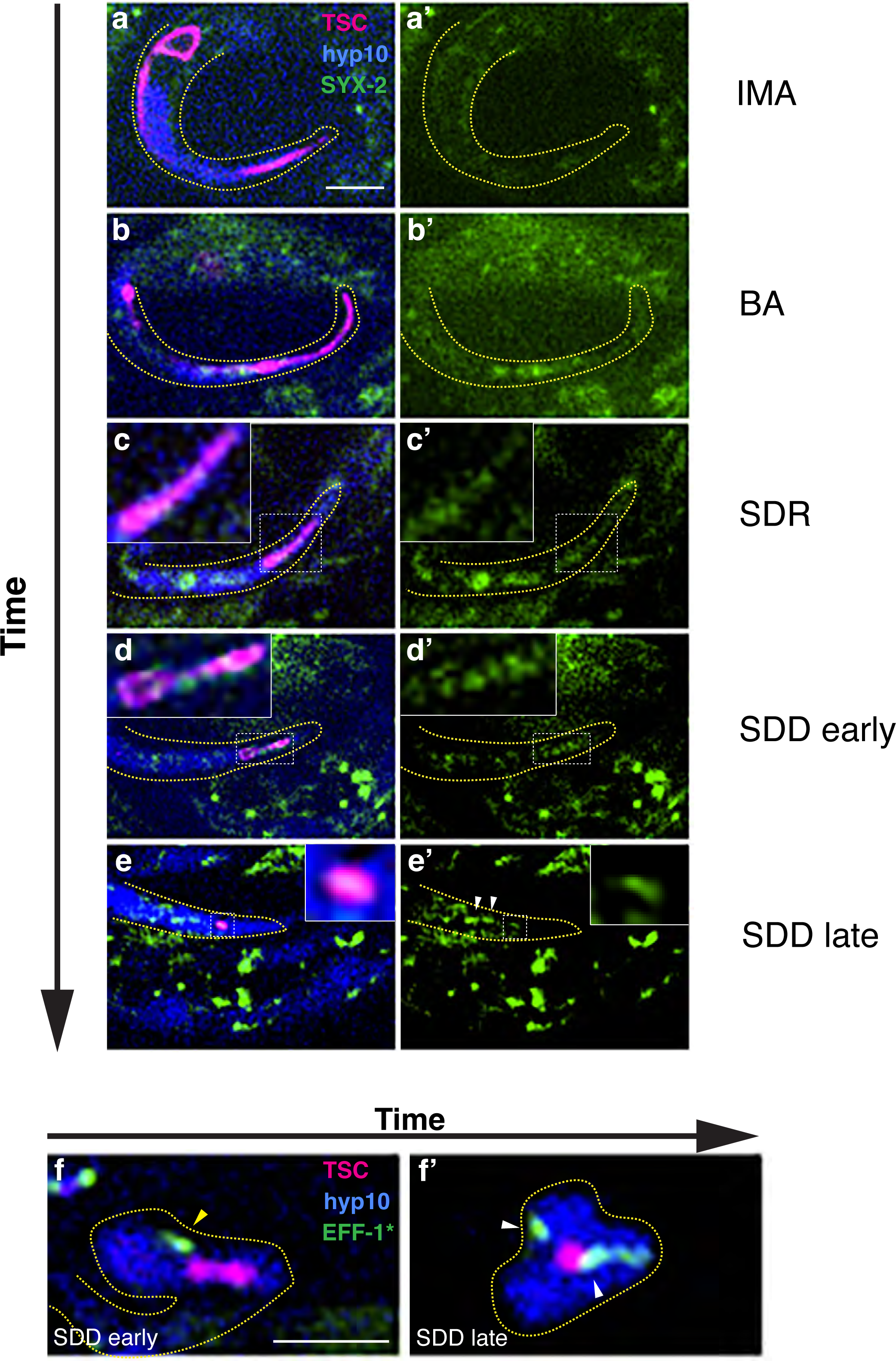
SYX-2/Syntaxin functions downstream of CDH-3/cadherin, YAP-1/YAP and EGL-44/TEAD. **(a-e’)** Tricolor labeling of GFP::SYX-2 (endogenously tagged), TSC (mKate2) and hyp10 (iBlueberry) across CCE stages. Representative localizations 5/5 for each CCE stage. (**f**) hyp10::EFF-1::GFP, TSC (mKate2) and hyp10 (iBlueberry) at the early soma-distal degrading (SDD) stage, yellow arrow showing EFF-1 on its way to distal TSC corpse, corresponding to (**d**) of GFP::SYX-2. (**f’**) hyp10::EFF-1::GFP, TSC (mKate2) and hyp10 (iBlueberry) at the late soma-distal degrading (SDD) stage, white arrows showing EFF-1 around distal TSC corpse in pseudopods, corresponding to (**e**) of GFP::SYX-2, white arrows showing SYX-2 not localizing around distal TSC corpse. Scale bar: 5μm.

Having identified the SDD stage of CCE as an important timepoint for SYX-2 function and the junction of the two connected distal process lobes an important site of function, we selected the SDD/bi-lobed distal stage (**Figure 4d, d’**) to test whether the CDH-3/YAP-1/EGL-44 axis may regulate SYX-2. Previous work has shown that SYX-2 physically associates EFF-1 binding at the EFF-1 C-terminus (41). Based on this and our present study we asked how SYX-2 may affect EFF-1. Interestingly, we found that at the bi-lobed distal process stage of the TSC, when SYX-2 centrally localized **(Figure 4d, d’**), the hyp10 EFF-1 appears to be approaching the distal TSC corpse **(Figure 4f**). In the following late stage when the distal process has reduced, while SYX-2 appears to be remote from the TSC **(Figure 4e, e’**), EFF-1 appears at near the TSC at the pseudopods **(Figure 4f’**). We therefore propose that SYX-2 aids in the recruitment of EFF-1 to the hyp10 phagocyte pseudopods, akin to its reported role in wound healing (41).

Moreover, while we found very discrete and specific signal in hyp10 around the TSC process in wild-type, this is localization pattern is not found in either *cdh-3* and *egl-44* mutants **(Figure 5b-d**). However, we did notice GFP signal more broadly in the hyp10. This suggests that CDH-3/YAP-1/EGL-44 do not affect SYX-2 transcription and that another factor that is important for SYX-2 localization is the transcriptional target. Our future studies will investigate potential downstream targets of the CDH-3/YAP-1/EGL-44 axis by drawing upon work in wound healing to explore the involvement of various SYX-2 regulators including RIC-4/SNAP, SEC-22/Synaptobrevin (56), and ESCRT components VPS-32.1 and VPS-4, and the involvement of actin polymerization (41). In our present study, to further ascertain whether SYX-2 indeed functions downstream of the newly identified CCE promoting CDH-3/YAP-1/EGL-44 axis, we overexpressed hyp10-specific *syx-2* in both *cdh-3* and *egl-44* mutants and found this to rescue the mutant defects (**Figure 5e, f**).

**Figure 5.**
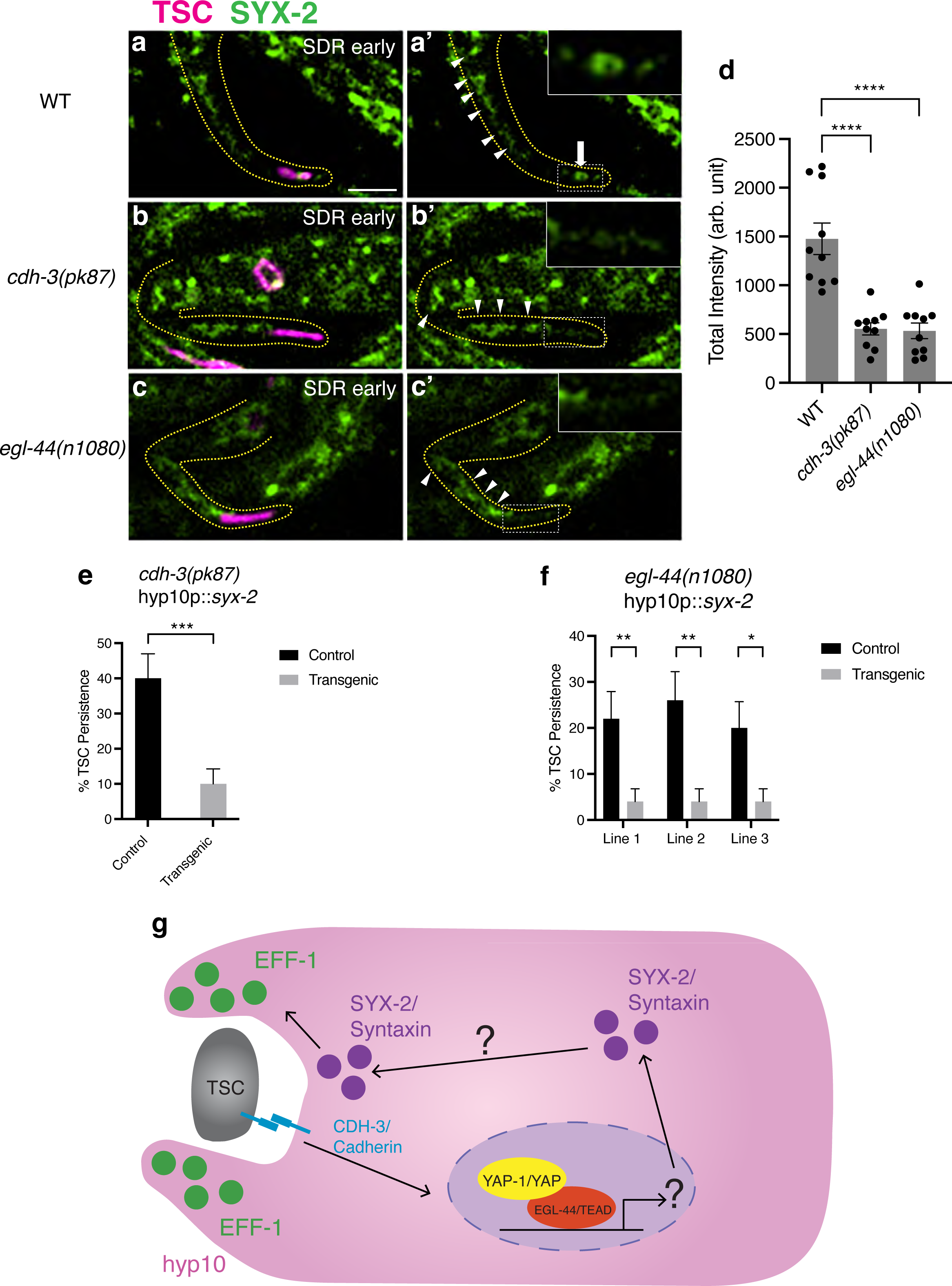
Endogenously-tagged (GFP) *syx-2* in wild-type(**a-a’**), white arrow showing SYX-2 accumulation around distal TSC corpse, *cdh-3(pk87)* (**b-b’**) and *egl-44(n1080)* (**c-c’**) backgrounds, white arrowheads showing SYX-2 in hyp10 not localizing around distal TSC corpse, and intensity quantifications (**d**). (**e**) Graph for TSC persistence for *syx-2* overexpression in *cdh- 3(pk87)* and *egl-44(n1080)* (f) mutants. (g) Cell-cell physical proximity changes between TSC and hyp10 are communicated promotes YAP-1 and EGL-44 function in positively regulating SYX-2. SYX-2/Syntaxin in turn promotes translocation of EFF-1/fusogen to the hyp10 phagocyte pseudopods, enabling phagosome closure. Scale bar: 5μm. For GFP quantification n=10. For over expression lines n>50. ns (not significant) *p* > 0.05, * *p* ≤ 0.05, ** *p* ≤ 0.01, *** *p* ≤ 0.001, **** *p* ≤ 0.0001.

Collectively, our data suggests that the SYX-2 localization is facilitated by YAP-1/YAP and EGL-44/TEAD as a response to CDH-3-mediated adhesion changes between the dying TSC and hyp10 phagocyte during CCE (**Figure 5g, Model**). We propose that localized SYX-2/Syntaxin may play a role in TSC process scission and/or hyp10 membrane autofusion, which in turn promotes translocation of EFF-1/fusogen to the hyp10 pseudopod membrane for phagosome closure.

## Discussion

Here we introduce a previously unreported regulatory pathway that promotes the role of the fusogen EFF-1 in phagosome sealing. Phagosome sealing is an otherwise poorly characterized step of phagocytosis, with EFF-1 identified as required for sealing of the nascent phagosome during the compartment-specific clearance of the TSC. We shed light on physical interaction via cell-cell adhesion as a possible initiation cue for phagocytosis. We report the unexpected observation that the hyp10 phagocytic cup appears to form around the TSC distal node well before the TSC has retracted and a reduced node has formed. This suggests that mechanical force between a dying cell and its phagocyte at the TSC distal node region may serve as a novel “eat-me” corpse recognition signal where the dying cell expands in a compartment-specific manner. We also present an open question as to what the transcriptional target of YAP-1 and EGL-44 is with respect to cell clearance. Given the impact of CDH-3/YAP-1/EGL-44 on the localization of SYX-2 protein specifically as opposed to its transcription, plausible candidates for transcriptional targets of our newly identified regulatory axis are proteins that have been shown in previous studies to regulate SYX-2 localization, including SNAP, Synaptobrevin, ESCRT components and regulators of actin (41, 56). We newly implicate a syntaxin in a new cell fusion role in the context of cell clearance. We also ask whether apparent sequential recruitment of SYX-2 and EFF-1 to the hyp10 pseudopods plays a specific role in the apparent scission of the TSC bi-lobed distal process to be internalized. It will also be interesting to explore whether cell-cell adhesion can impact other fusogens, such as *C. elegans* AFF-1 fusogen, shown to act as an exoplasmic fusogen during tube elongation (57). Finally, we raise the intriguing point that the cell biological basis of wound healing and phagocytosis share mechanistic overlap, both involving controlled membrane fusion events.

## Materials and Methods

### C. elegans methods

(a) *C. elegans* strains were cultured using standard methods on *E. coli* OP50 and grown at 20°C. Wild-type animals were the Bristol N2 subspecies. For most TSC experiments, one of two integrated reporters were used: *nsIs435* or *nsIs685.* For hyp10 experiments, *nsIs836* was used. Integration of extrachromosomal arrays was performed using UV and trioxsalen (T2137, Sigma). Animals were scored at 20°C.

### Imaging

Images were collected on a Nikon TI2-E Inverted microscope using a CFI60 Plan Apochromat Lambda 60x Oil Immersion Objective Lens, N.A. 1.4 (Nikon) and a Yokogawa W1 Dual Cam Spinning Disk Confocal. Images were acquired using NIS-Elements Advanced Research Package. For still embryo imaging, embryos were anesthetized using 0.5 M sodium azide. Larvae were paralyzed with 10mM sodium azide.

### Quantifications

For CCE defects, TSC death defects were scored at the L1 stage. Animals were mounted on slides on 2% agarose-M9 pads, paralyzed with 10mM sodium azide, and examined on a Zeiss Axio-Scope A1. The persisting TSC was identified by fluorescence based on its location and morphology.

Quantification of Fluorescence Intensity: Sum intensity projections of fluorescent reporters in relevant cell regions (hyp10) were generated by following TSC membrane signal in ImageJ software, and GFP intensity was measured. Corrected Total Cell Fluorescence (CTCF) was calculated using Microsoft Excel and graphed using GraphPad Prism. Statistical analysis: unpaired two-tailed t-test for comparison between wild-type and mutant animals.

### Worm strains used in this study

LGII-*egl-44 (n1080)*, *eff-1(ns634)*

LGIII-*cdh-3 (pk87), cdh-3 (pk77), cdh-3(mcc39)*

LGX- *yap-1*(ys38*), syx-2(mcc35), syx-2(mcc40)*

### Plasmids and Transgenics

Plasmids were generated via Gibson cloning. Primer sequences and information on the construction of plasmids used in this study are provided in (**Supplemental Table 1**). The full list of transgenes is described in (**Supplemental Table 2**). The full length or fragment of the *aff-1* promoter was used to label the TSC. The *eff-1* promoter was used to label hyp10.

### CRISPR Cas9 genome editing

The alleles of *syx-2(mcc35)* and *syx-2(mcc40)* were made by introducing GFP just upstream of the start of the *syx-2* gene via CRISPR/Cas9 to generally endogenously N-terminally tagged GFP::SYX-2. Mutants were generated using a co-injection strategy (58). Guide crRNA, repair single-stranded DNA oligos, tracrRNA, and buffers were ordered from IDT. Guide crRNA used to generate *syx-2(mcc35)* and *syx-2(mcc40)* was 5’-TGAAGAAGGACAAAGTCAAT -3’.

The allele of *cdh-3(mcc39)* was made by introducing GFP just upstream of the stop of the *cdh-3* gene via CRISPR/Cas9 to generally endogenously N-terminally tagged CDH-3. Mutants were generated using a co-injection strategy (58). Guide crRNA, repair single-stranded DNA oligos, tracrRNA, and buffers were ordered from IDT. Guide crRNA used to generate *cdh-3(mcc39)* was 5’-CATTTTCGATTCATTTTTAC-3’.

### Statistics

Sample sizes and statistics were based on previous studies of CCE and the TSC (44). Independent transgenic lines were treated as independent experiments. An unpaired two-tailed *t-* test was used for all persisting TSC quantifications (GraphPad Prism). For all figures, mean ± standard error of the mean (s.e.m.) is represented.

### Data Availability Statement

Strains and plasmids are available upon request. The authors affirm that all data necessary for confirming the conclusions of the article are present within the article, figures, and tables.

### Conflict of Interest Statement

There is no conflict of interest to disclose.

## Acknowledgments

AW and PG designed the experiments and wrote the manuscript. AW and AE performed the experiments. GC helped generate strains. AW, AE and PG analyzed the data. We thank members of the Ghose lab for comments on the manuscript. Some strains were provided by the CGC, which is funded by NIH Office of Research Infrastructure Programs (P40 OD010440). We thank Dr. Junho Lee for providing strains.

## Funding

PG is a Cancer Prevention Research Institute of Texas (CPRIT) Scholar in Cancer Research (RR100091) and is also funded by a National Institutes of Health-National Institute of General Medical Sciences Maximizing Investigators’ Research Award (MIRA) (R35GM142489).

## Supplemental Figures

**Figure S1.**
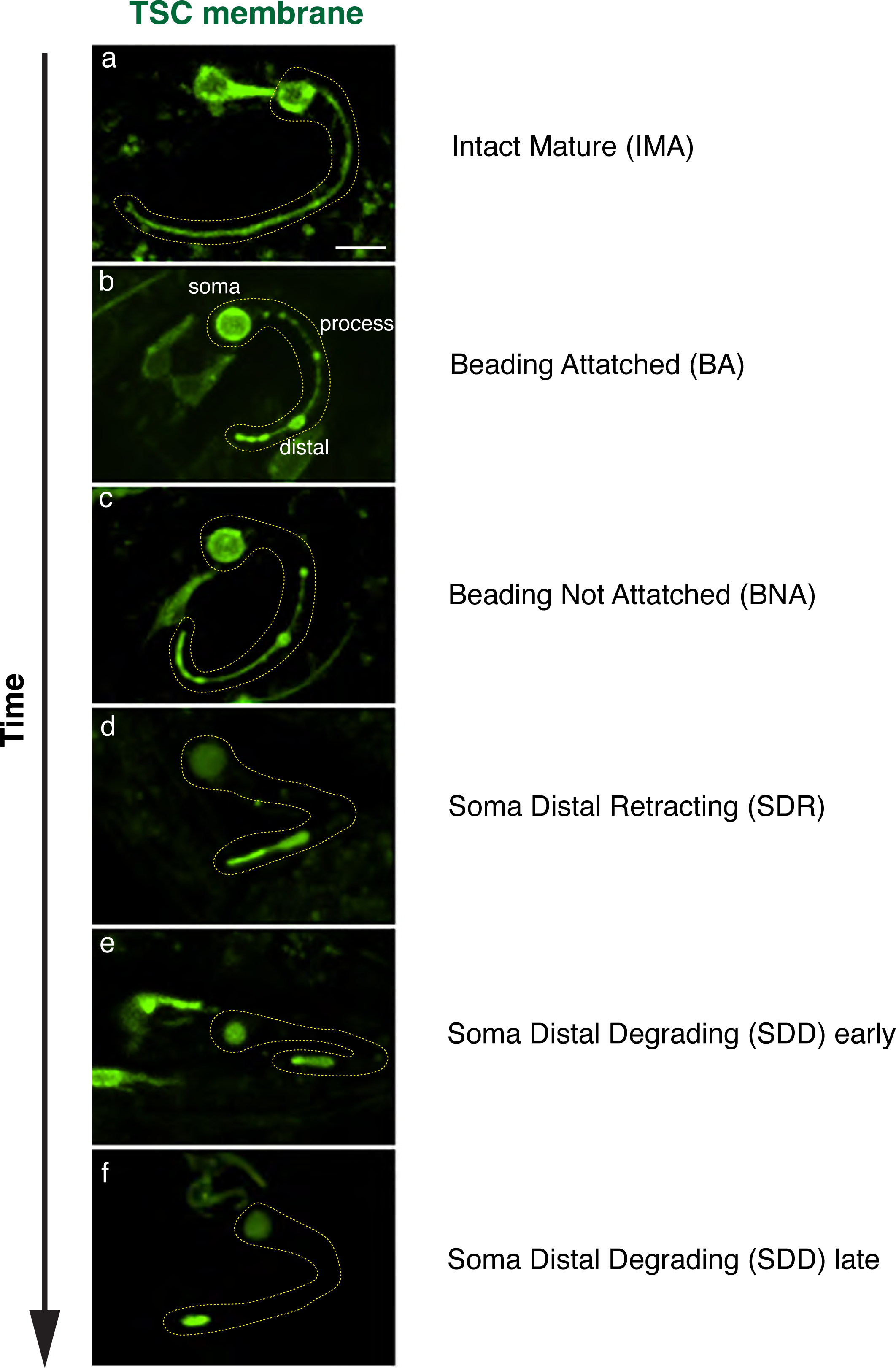
CCE stages and designations. TSC membrane reporter labeled in green. Wild-type CCE stages shown (a-f). (a) intact mature, (b) beading attached, (c) beading not attached, (d) soma distal retracting, (e) soma distal degrading early, (f) soma distal degrading late.

**Figure S2.**
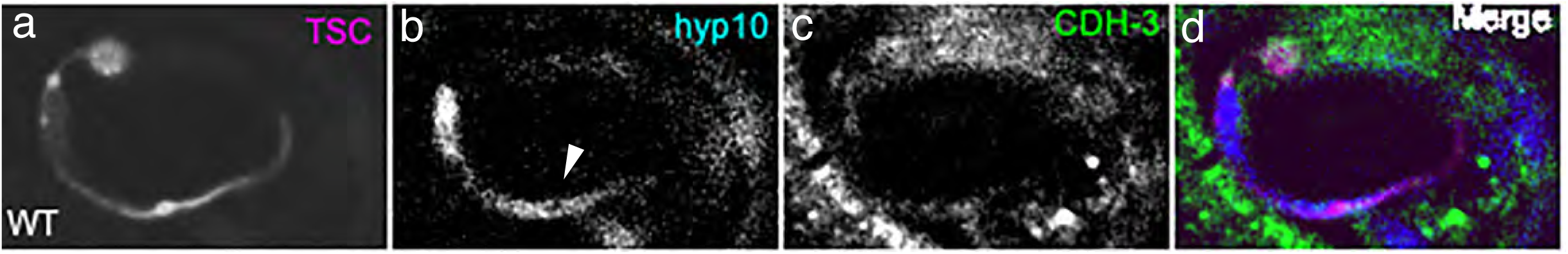
Hyp10 phagocytic cup formation (white arrowhead) around newly formed distal node during CCE beading stage.

**Figure S3.**
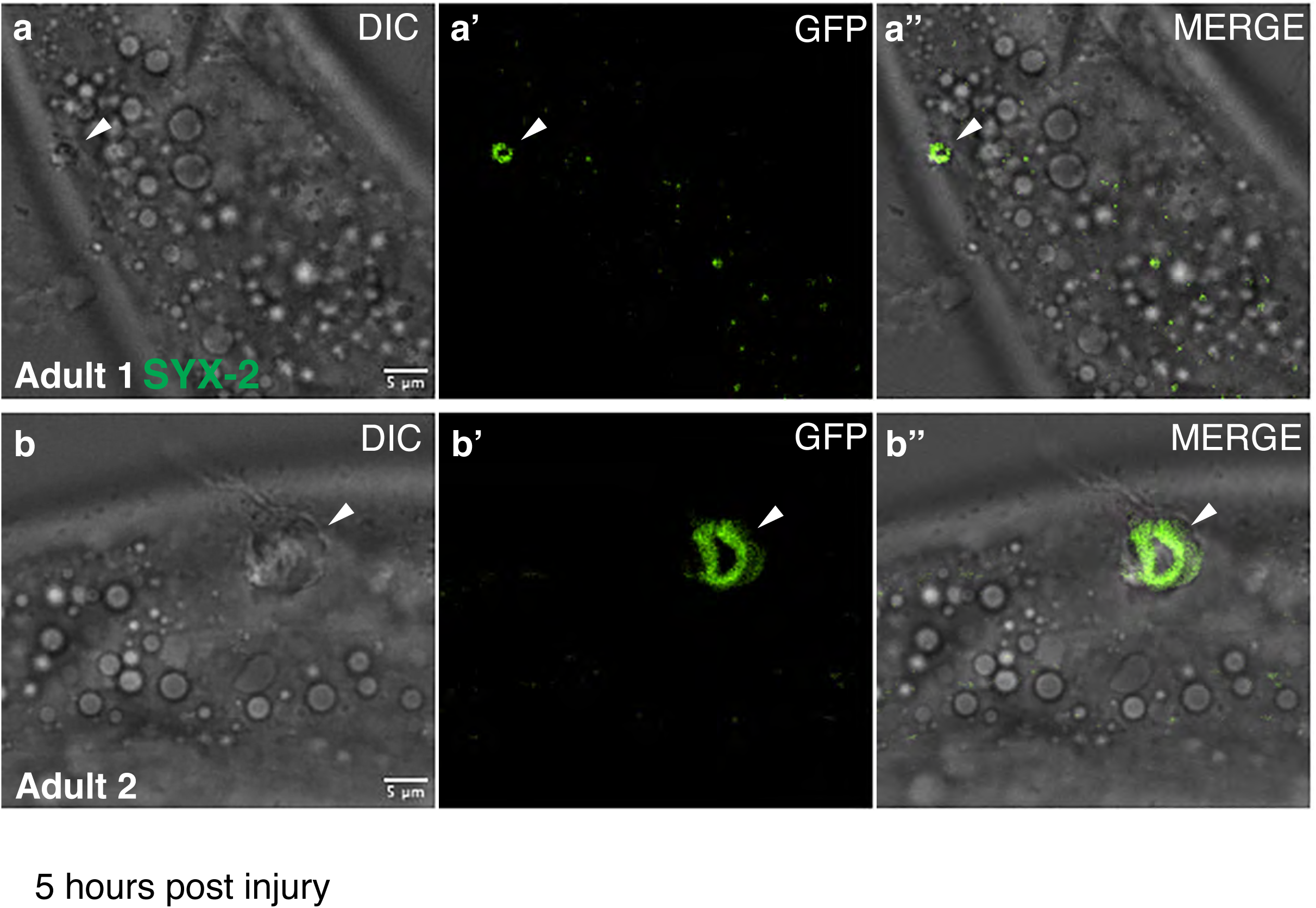
Validation of CRISPR Cas9-generated GFP::SYX-2 via injury induced wound healing response at five hours post injury per published protocol (41). Wound marked with white arrowhead.

## Supplemental Movies

Movies are Z-series of images in main figures.

M1. CDH-3 localizes to both the TSC and hyp10 from the TSC beading stage of CCE (BA).

M2. CDH-3 localizes to both the TSC and hyp10 from the TSC soma-distal retracting stage of CCE (SDR).

M3. CDH-3 localizes to both the TSC and hyp10 from the TSC soma-distal degrading stage of CCE (SDD).

M4. TSC internalization failure in *cdh-3(pk87)* mutants.

M5. TSC internalization failure in *eff-1(ns634)* mutants.

M6. EFF-1 localization in wild-type.

M7. EFF-1 localization defect *cdh-3(pk87)* mutants (L1)-no signal.

M8. EFF-1 localization defect *cdh-3(pk87)* mutants (L1)-weak signal.

M9. TSC internalization failure in *yap-1(ys38)* mutants.

M10. TSC internalization failure in *egl-44(n1080)* mutants.

M11. SYX-2 localization at CCE SDR.

M12. SYX-2 localization at CCE SDD early.

M13. SYX-2 localization at CCE SDD late.

M14. EFF-1 of hyp10 appearing to approach the TSC distal remnant at soma-distal degrading early stage of CCE (SDD).

M15. EFF-1 at hyp10 pseudopods soma-distal degrading late stage of CCE (SDD).

M16. SYX-2 localization in wild-type.

M17. SYX-2 localization defect *cdh-3(pk87)* mutants (L1).

M18. SYX-2 localization defect *egl-44(n1080)* mutants (L1).

## Notes

### Competing Interest Statement

The authors have declared no competing interest.

